# Live vibratome sectioning of human lung and cerebral organoids for infection studies with respiratory and neurotropic viruses

**DOI:** 10.64898/2026.11.23.768492

**Authors:** Simon T. Herrmann, Wing Hang Ip, Luca D. Bertzbach, Thorsten Müller, Esther Schnettler, Stephanie Pfaender

## Abstract

We present a robust, reproducible protocol for generating live, thin (300 µm) vibratome sections from human lung organoids and human cerebral organoids. The method preserves the native three-dimensional tissue architecture, cellular heterogeneity, and viability for at least 7 days in culture, providing a physiologically relevant platform for investigating respiratory and neurotropic virus infection. Following infection, samples can be fixed, cryoprotected, and embedded for downstream cryosectioning. These organoid sections support efficient analyses of viral entry, replication, and host responses. The technique is readily compatible with immunofluorescence, RNA *in situ* hybridization, spatial transcriptomics, live-cell imaging, and a wide range of advanced molecular and cellular assays. On average, five sections can be obtained from a single organoid, substantially increasing experimental throughput while reducing costs and biological variability. By combining the physiological complexity of three-dimensional organoid systems with the accessibility and scalability of tissue-section-based assays, this platform bridges a critical gap in respiratory and neuronal disease modeling. The protocol provides a versatile and high-resolution framework for studying respiratory and neurotropic viral pathogenesis, host– pathogen interactions, and therapeutic interventions in human lung as well as brain tissue.

## 1. Introduction

Human lung organoids (HLOs) recapitulate key structural and functional features of the human airway epithelium, including upper airway cell types such as ciliated, secretory, and basal progenitor cells, as well as lower airway/alveolar cell types including alveolar type I and type II (AT1/AT2) cells, making them valuable models for studying respiratory viral infections such as severe acute respiratory syndrome coronavirus 2 (SARS-CoV-2), the causative agent of COVID-19 (1-3). Their ability to self-organize into 3D structures preserving cellular heterogeneity and polarized epithelium has positioned them as a transformative tool in respiratory research. Similarly, air-liquid interface (ALI) cultures and precision-cut lung slices (PCLS) have emerged as complementary platforms: ALIs provide a pseudostratified, polarized two-dimensional epithelium for studying viral entry and immune responses, while PCLS preserve the native tissue architecture, immune cell infiltration, and stromal microenvironment, enabling studies of tissue-level pathology (4-7).

Likewise, human cerebral organoids (HCOs) have emerged as powerful *in vitro* models for studying neurodevelopmental processes and neurotropic viral infections. These 3D structures recapitulate key features of the developing human brain, including layered cortical organization, diverse neuronal subtypes, and functional neural networks (8, 9). They provide a human-relevant platform for investigating host-pathogen interactions in the central nervous system, particularly for viruses with neuroinvasive potential, such as West Nile virus (WNV) (10).

However, the 3D architecture of organoids limits access to internal cell populations and hinders uniform viral exposure and imaging (1). This diffusion barrier restricts the ability to study viral tropism, replication dynamics, and host responses in subepithelial and stromal compartments. To overcome these limitations, we used vibratome sectioning of cerebral organoids, a method first reported by Giandomenico et al. (11). This approach generates thin, viable organoid slices that preserve native cellular composition and spatial organization while providing direct access to the epithelial surface and underlying stromal regions. Janssens et al. previously infected 500-µm-thick sectioned cerebral organoids with Zika virus, but infection rates were poor compared to 2D dissociation approaches (12). Another alternative approach for viral infection of organoids is microinjection (13, 14), although this technique can be technically challenging under high containment conditions. This protocol describes the embedding of HLOs and HCOs in low-melting-point agarose (LMP agarose), followed by precision sectioning using a vibratome. The resulting live sections maintain viability and morphological integrity, enabling infection with respiratory or neurotropic viruses, exemplified by SARS-CoV-2 and WNV, respectively, and downstream analysis of viral replication, host immune responses, and tissue-level pathology. To our knowledge, this is the first report of WNV infection of sectioned cerebral organoids and the first detailed protocol for SARS-CoV-2 infection of sectioned lung organoids, extending the previously reported use of sliced lung organoids for antiviral testing (15). The method is optimized to maximize reproducibility, minimize tissue damage, and ensure compatibility with a wide range of downstream applications. Importantly, multiple viable sections can be generated from a single organoid, allowing different experimental conditions, such as treated versus untreated or infected versus mock-infected samples, to be evaluated within the same biological specimen. This approach substantially reduces biological variance arising from organoid-to-organoid heterogeneity, improves statistical robustness, and enables more sensitive comparisons between experimental groups. By combining the genetic tractability and scalability of human induced pluripotent stem cell (hiPSC)-derived organoids with the spatial resolution and accessibility of precision-cut tissue slices, vibratome slices offer a powerful, high-throughput platform for dissecting host-pathogen interactions in human lung or cerebral tissue.

## 2. Materials

### 2.1. Organoids and reagents

- STEMdiff Branching Lung Organoid Kit (Stemcell Technologies) resulting in human lung organoids (HLOs), diameter ∼1 mm
- STEMdiff Cerebral Organoid Kit (Stemcell Technologies) resulting in human cerebral organoids (HCOs), diameter ∼1 mm
- hiPSCs
- UltraPure Low Melting Point (LMP) Agarose (Invitrogen)
- DMEM/F12
- Dulbecco’s Phosphate-Buffered Saline (DPBS)
- BLO-B medium (HLO cultivation media; Stemcell Technologies)
- Maturation medium (HCO cultivation media; Stemcell Technologies)
- Matrigel (for re-embedding if needed)
- Anti-adherence rinsing solution (Stemcell Technologies)
- Superglue
- Phosphate-buffered saline (PBS, pH 7.4))
- 4% Paraformaldehyde (PFA) in PBS
- 30% (w/v) sucrose in PBS
- Tissue-Tek OCT compound
- Purple loading dye
- Superfrost plus slides

### 2.2. Equipment

- Vibratome (Leica VT 1200S or equivalent)
- Heating block
- Ice bucket with ice
- Sterile 1 cm^3^ silicone mold (EtOH or heat sterilized)
- Scalpel
- Fine tweezers
- Glass slides
- Dry ice
- Cryostat
- -80°C freezer
- Fine paintbrush
- Stainless steel spatula
- P200 and P1000 pipettes with cut tips (sterile, cut them with clean scissors)
- 1.5 mL microcentrifuge tubes
- 48-well tissue culture plates
- Petri dishes (60 mm)
- Microscope (phase-contrast or fluorescence, for section inspection)

## 3. Methods

Before you start vibratome/live sectioning of organoids: HLOs and HCOs are generated from hiPSCs using defined differentiation protocols and cultured in 3D Matrigel matrices. For vibratome sectioning, select organoids with a diameter of at least 1 mm (approximately 80 days of differentiation) to ensure sufficient organoid volume, structural integrity, and consistent section quality. Smaller organoids (<500 µm) are prone to fragmentation and may not yield usable sections.

### 3.1. Preparation of low-melting-point agarose

1. Prepare a 3% (w/v) solution of UltraPure LMP agarose in DMEM/F12 under sterile conditions.
2. Aliquot 1.2 mL portions and store at 4°C until use.
3. Prior to embedding, liquefy agarose at ca. 80°C using a heating block (**see Note 1**).
4. Reduce the temperature to 40°C and maintain until embedding.

### 3.2. Isolation of HLOs from Matrigel

1. Collect HLOs cultured in a Matrigel-sandwich configuration. For reproducibility, we used the above-mentioned Stemcell kit according to the manufacturer’s instructions (**see note 2**)
2. Remove culture medium and invert the transwell insert into a petri dish using tweezers.
3. Cut the insert membrane at the edge with a scalpel and lift it off carefully.
4. If the Matrigel-sandwich does not detach spontaneously, gently wash it out using DPBS (**see Note 3**).
5. Using a scalpel, carefully slice the Matrigel to separate individual organoids and remove excess Matrigel (Fig 1).
6. Transfer organoids into cultivation medium using a cut P1000 pipette tip until embedding.

**Figure 1.**
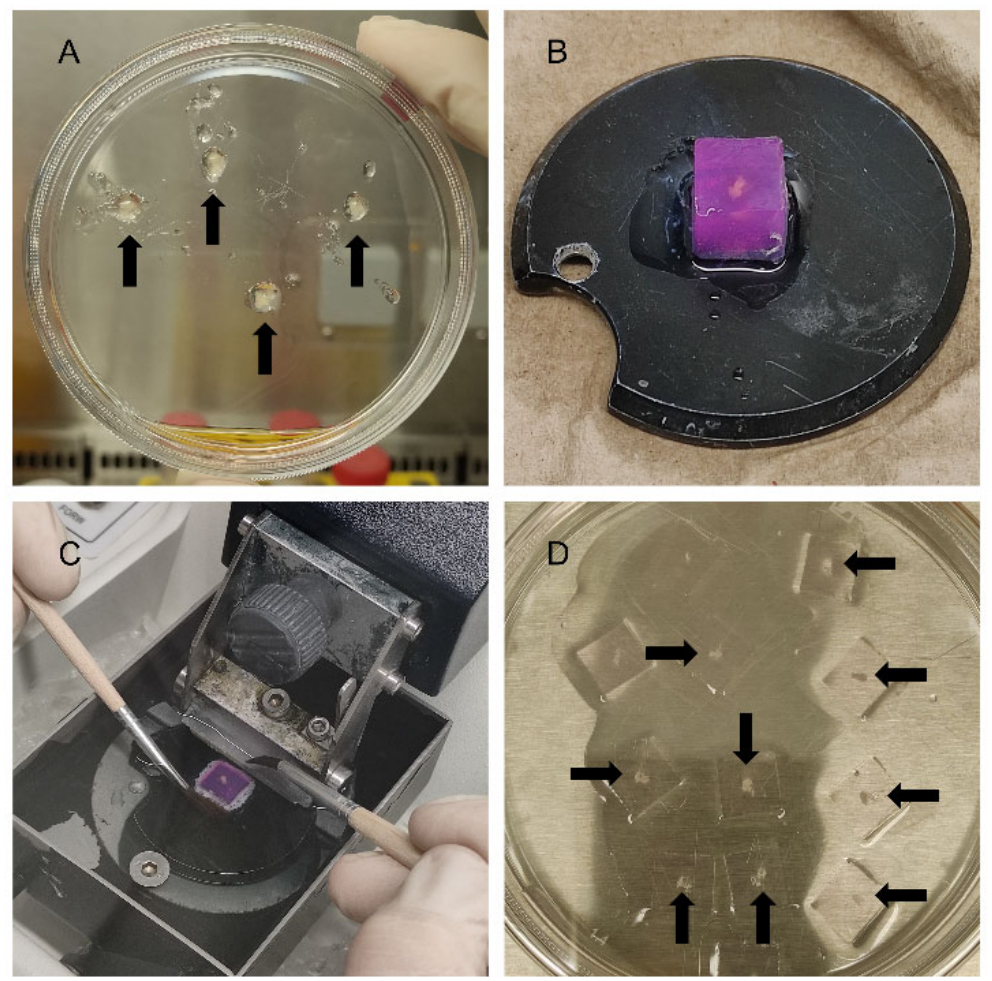
Workflow of live vibratome sectioning of human organoids. Schematic representation of key steps in the preparation of thin (300 µm) vibratome sections from organoids. (**A**) Manual dissection of organoids from residual Matrigel using a scalpel; individual organoids are separated and washed in DPBS. (**B**) Embedding of a single organoid in a 3% low-melting-point agarose cube, which is mounted onto the vibratome specimen holder using superglue in the desired orientation. (**C**) Vibratome sectioning: the oscillating blade cuts through the agarose-embedded organoid. A fine paintbrush is used to gently lift and collect freshly cut sections. (**D**) Collection of organoid sections in a petri dish filled with ice-cold DPBS. Arrows indicate 10 intact sections still attached to the agarose matrix.

### 3.3. Embedding of organoids in agarose

1. Transfer organoids into a 1.5 mL microcentrifuge tube using a cut P1000 pipette tip, remove cultivation media as much as possible.
2. Fill a sterile 1 cm^3^ silicone mold halfway with liquefied 3% LMP agarose.
3. Add a small volume of agarose to the organoids using a cut P1000 pipette tip and transfer the organoid-agarose mixture into the mold (**see Notes 4 and 5**).
4. Fill the mold completely with agarose.
5. Gently stir the organoid within the agarose to ensure uniform stiffness and position it centrally using a P200 pipette tip.
6. Place the mold on ice for approximately 5 minutes to allow complete solidification (Table 1).
7. Carefully remove the hardened agarose block from the mold using a spatula or tweezers.

**Table 1.** Troubleshooting guide.

| **Problem** | **Likely cause** | **Solution** |
|---|---|---|
| **Tissue tearing during sectioning** | Blade too fast or frequency too high | Reduce speed to 4–5; lower frequency to 5–6 |
| **Organoid displaced from center** | Agarose solidified before organoid added | Work faster; pre-warm agarose to 40°C only |
| **Agarose not solidifying** | Agarose cooled below 35°C | Warm to 40°C before use |
| **Poor section adhesion to plate** | Plate not properly coated | Re-coat with anti-adherence solution |
| **Low viral signal** | Low MOI or degraded virus | Use fresh stock, increase PFU/section |
| **Poor section adhesion** | Insufficient OCT freezing | Freeze sample 5 min before sectioning |
| **Ice crystals in sections** | No cryoprotection | Use 30% sucrose; freeze slowly |
| **Weak immunofluorescence signal** | Over-fixation or poor blocking | Fix 24 h; block 1 h with 5% milk/TBST |
| **High immunofluorescent background** | Antibody too concentrated | Titrate antibodies; wash 5× with TBST |
| **High background in staining** | Residual agarose | Gently scrape with scalpel or wait 24 h |
| **Sample dislodged** | Excess sucrose or loose Parafilm | Blot excess fluid; wrap slide tightly |

### 3.4. Vibratome sectioning

1. Mount the agarose block onto the vibratome specimen holder using superglue, ensuring the desired orientation for sectioning. Allow the adhesive to cure for approximately 2 minutes until fully set before proceeding.
2. Place the specimen holder into the vibratome chamber filled with ice-cold DPBS and surround the chamber with ice.
3. Section the embedded organoids at a thickness of 300 µm (**see Note 6**).
  ∘ Blade frequency: 7
  ∘ Cutting speed: 6
4. Collect sections using a fine paintbrush and transfer them into a petri dish. Alternatively, sections may be transferred directly into a 24 or 48-well plate (previously coated with anti-adherence rinsing solution and washed once with DMEM/F12) containing culture medium (Figure 1).

### 3.5. (Optional) removal of agarose and culture of sections

1. After sectioning, carefully remove excess agarose from each section using a scalpel (**see Note 7**).
2. Collect organoid sections with a cut P1000 pipette tip in cultivation medium.
3. Transfer sections into a 24 or 48-well plate previously coated with anti-adherence rinsing solution and washed once with DMEM/F12.
4. Culture sectioned organoids at 37°C, 5% CO_2_, in a humidified incubator. Sample quality can be checked by light microscopy (Figure 2).

**Figure 2.**
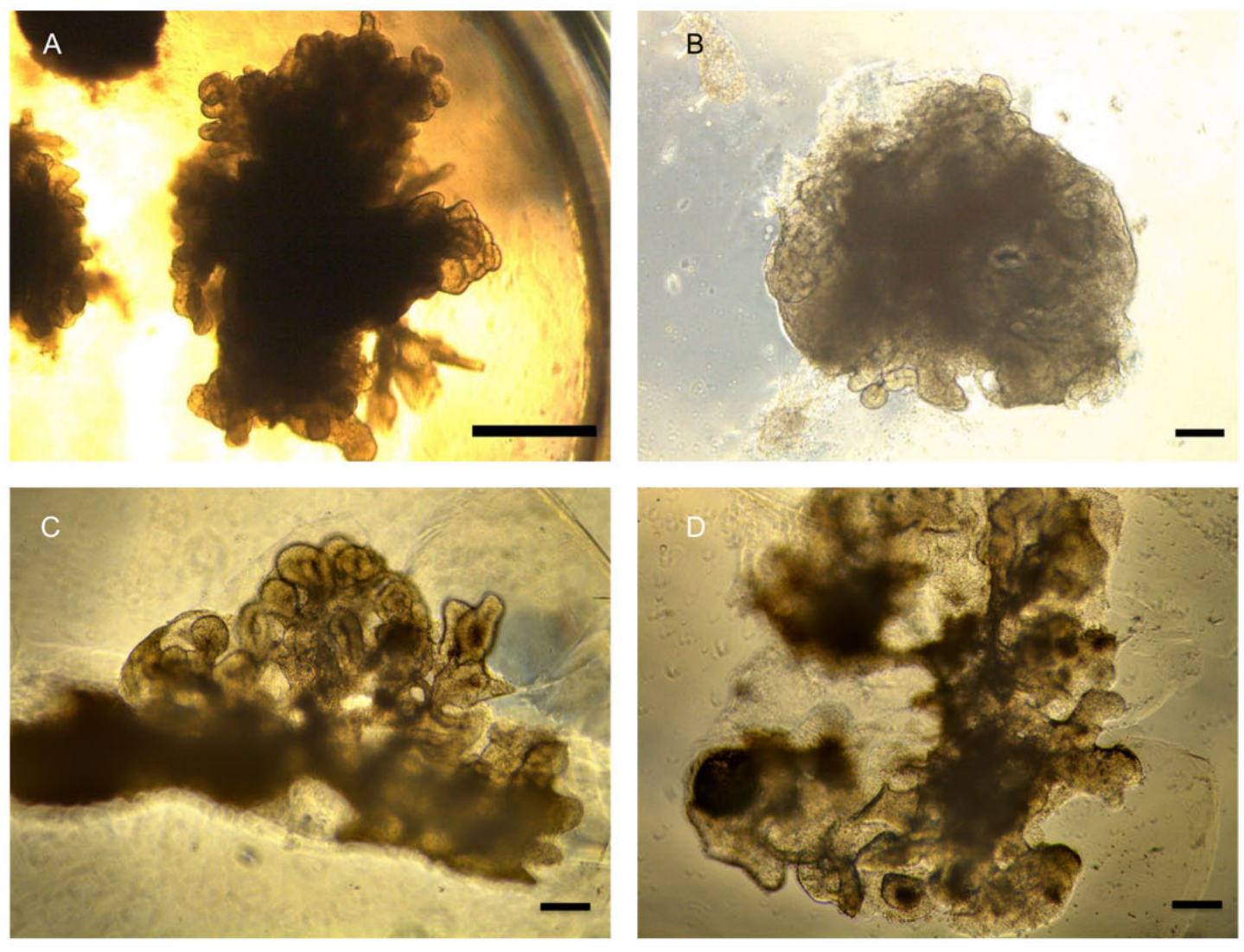
Morphological comparison of HLOs before and after vibratome sectioning. Bright-field microscopy images illustrating the structural changes during the sectioning process. (**A**) Intact HLO embedded in a Matrigel-sandwich prior to sectioning; scale bar: 1000 µm. (**B-D**) Representative 300 µm vibratome sections of the same HLO, showing preserved 3D architecture, cellular heterogeneity, and epithelial organization. Scale bars: 200 µm.

### 3.6. Infection of vibratome sections

1. For viral infection (e.g., SARS-CoV-2, WNV; Figs. 3 and 4), prepare the inoculum in culture medium containing 35,000 PFU of virus per section. We recommend using a 48-well plate pre-coated with anti-adherence solution, placing one vibratome section per well in a final culture volume of 200 - 400 µL (**see Notes 8-12**).
2. Incubate the samples at 37°C, 5% CO_2_, in a humidified incubator for 2 h to allow viral attachment and entry.
3. After 2 h, gently wash the sections three times with 400 μL of pre-warmed PBS (37°C) to remove unbound virus.
4. Replace the wash buffer with 200 - 400 μL of fresh cultivation medium.
5. Culture the samples at 37°C, 5% CO_2_, in a humidified incubator.
6. (Optional) replace the medium daily with fresh cultivation medium.
7. Collect the supernatant from each well at, e.g., 24, 48, and 72 h post-infection for downstream analysis, such as end-point dilution assays.
8. At 72 h post-infection, wash the sections three times with 200 - 400 μL of PBS.
9. Proceed immediately to fixation (**see Note 12**).

**Figure 3.**
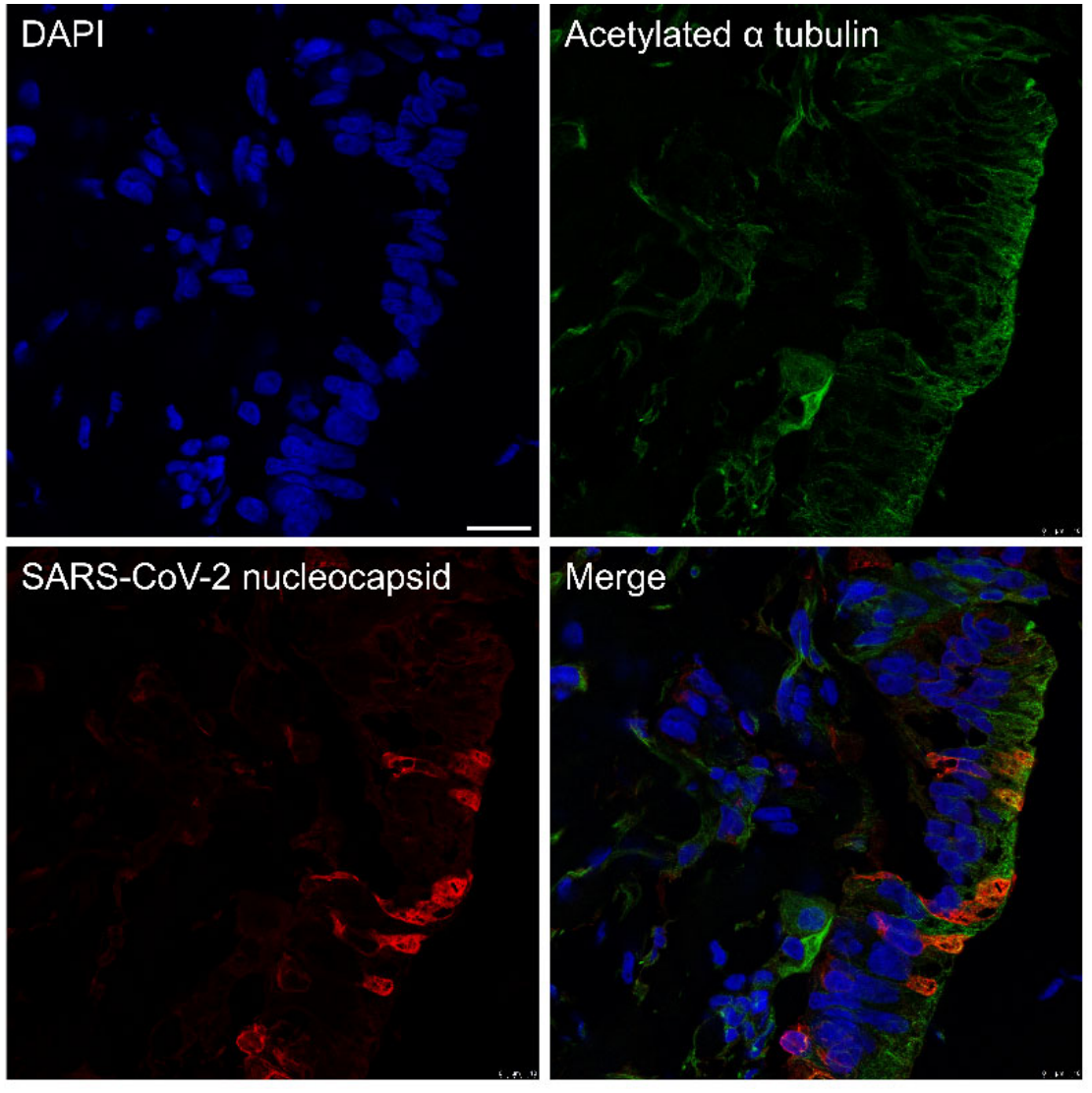
SARS-CoV-2 infection targets ciliated epithelial cells in HLO vibratome sections. Immunofluorescence analysis of a cryosection from a vibratome-sectioned HLO infected with SARS-CoV-2. DAPI: nuclear counterstain. Acetylated α-tubulin (green; Sc-23950, Santa Cruz): marker for ciliated cells. Nucleocapsid protein (N, red; MA5-36086, Invitrogen): SARS-CoV-2 antigen. Merged image showing co-localization of SARS-CoV-2 N protein (red) and acetylated α-tubulin (green) in ciliated cells (yellow signal). Scale bar: 20 μm. Image captured at 100× magnification.

**Figure 4.**
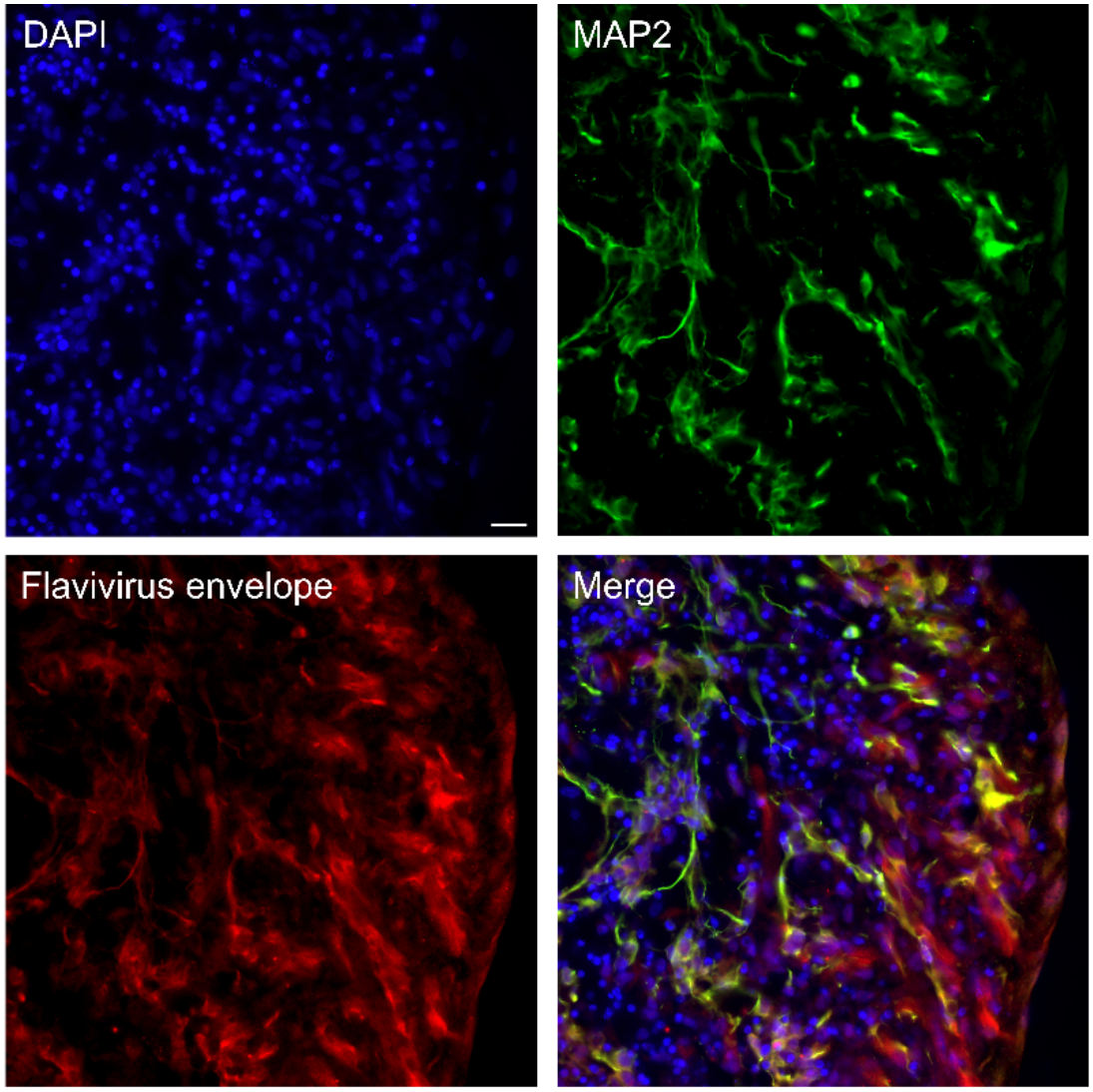
WNV infection of HCO vibratome sections. Immunofluorescence analysis of cryosections from a vibratome-sectioned HCO infected with WNV. DAPI (blue) counterstains nuclei. MAP2 (green; ab5392, Abcam) labels mature neurons (somata and dendrites). WNV envelope protein (E, red; GTX132052, GeneTex) marks WNV-infected cells. Scale bar, 20 µm; magnification, 60×.

### 3.7. Fixation and embedding of vibratome-sectioned virus-infected samples

1. Transfer vibratome-sectioned organoids from the 48-well plate to a sterile 1.5 mL microcentrifuge tube using a cut P1000 pipette tip.
2. Add 1 mL of 4% PFA in PBS to fix the samples.
3. Incubate at 4°C for 24 h to ensure complete fixation (**see Note 13**).
4. After fixation, carefully remove the PFA solution using a pipette.
5. Wash the fixed samples three times with 1 mL of cold PBS (4°C), using gentle pipetting to avoid dislodging the tissue.
6. Incubate the samples in 30% (w/v) sucrose in PBS at 4°C overnight for cryoprotection.
7. After cryoprotection, replace the sucrose solution with OCT compound/sucrose mixture (1:1 v/v) and proceed immediately to embedding.
8. Fill a 1 cm^3^ silicone mold halfway with pre-chilled OCT compound/sucrose mixture (1:1 v/v) and place on dry ice.
9. Once the bottom layer is partially frozen, transfer the sample using a cut P1000 pipette tip into the mold.
10. (Optional) place one drop of purple loading dye at one corner of the mold to mark the sample orientation within the frozen OCT compound/sucrose block. This facilitates later cryosectioning by enabling reliable identification of the sample position.
11. Carefully fill the mold completely with OCT compound/sucrose mixture.
12. Snap-freeze the mold on dry ice for 2-3 min, then transfer to -80°C for at least 12 h to ensure complete solidification.

### 3.8. Cryosectioning

1. Equilibrate the cryostat to -25°C for HLOs, -20°C for HCOs, at least 30 min prior to sectioning (**see Note 14**).
2. Place a thin layer of pre-chilled OCT compound (or equivalent) onto the cryostat holder or stage.
3. Allow the OCT layer to freeze completely in the cold chamber (typically 2-3 min; **see Note 15**).
4. Remove the frozen sample block from the silicone mold:
  ∘ Gently push the block from the bottom using a sterile spatula.
  ∘ The flexible, non-adhesive silicone mold facilitates easy release without damaging the sample.
5. Apply a small drop of OCT compound onto the pre-frozen OCT layer on the cryostat specimen holder, and position the frozen sample block on top.
6. If necessary, add a small drop of OCT compound around the block to secure it.
7. Use a fine tool to gently adjust the sample’s position for optimal orientation.
8. Return the holder to the cryostat chamber and allow the OCT to freeze for at least 10 min to ensure firm adhesion (Table 1).
9. Install the cryostat holder into the cutting position.
10. Begin sectioning at a thickness of 50 µm to trim the surrounding OCT and expose the organoid.
11. Once the organoid is visible, reduce the section thickness to 16 µm (**see Note 16**).
12. Mount 3-4 consecutive sections onto a single Superfrost Plus slide (or equivalent charged slide).
13. Allow the slide to air-dry for 5-10 min at room temperature before storing.
14. Store samples at -80° C until immunofluorescence staining (Table 1).

## 4. Notes

1. LMP agarose solidifies rapidly at room temperature. Work quickly after liquefaction to prevent premature gelation. For embedding, the agarose should be maintained at approximately 40 °C, as higher temperatures may compromise organoid viability.
2. HLOs are cultured in a Matrigel-sandwich. The Matrigel must be removed as good as possible prior to agarose embedding to ensure uniform and appropriate embedding for successful vibratome sectioning. In contrast, HCOs are typically embedded in Matrigel but grow outward, leaving only minimal residual Matrigel at the surface. In our hands, this small amount usually does not interfere with agarose embedding or sectioning and does not require removal.
3. Avoid prolonged exposure to DPBS or mechanical stress to preserve viability.
4. The organoid must be centrally embedded to ensure uniform sectioning. Avoid air bubbles during casting. Embedding should be performed in a single continuous pour (“single-step casting”) to ensure a homogeneous agarose matrix. If the agarose has already begun to solidify before the organoid is introduced, proper integration will not occur, resulting in an inhomogeneous block. This can critically impair downstream sectioning, as non-uniform agarose blocks are mechanically unstable and prone to fracture during vibratome slicing.
5. Most critical step is the embedding of the organoid in the agarose!
  - Melt the agarose at high temperature, then cool it to approximately 40°C. At this temperature, the agarose remains liquid while avoiding damaging of the sample.
  - It is recommended to practice this step using a substitute sample (e.g., an “organoid dummy”) before working with actual specimens.
  - If embedding is unsuccessful, carefully remove as much agarose as possible from the organoid and repeat the procedure.
6. Optimize blade frequency and speed empirically for organoid integrity. Higher frequencies may cause tissue tearing (Table 1).
7. Agarose may be left in place; it typically detaches within 24 - 48 hours. However, removal improves cell access and imaging.
8. Adhere to applicable national biosafety regulations and institutional regulations with appropriate containment and PPE.
9. Use only validated viral stocks with known titers (e.g., determined by plaque assay or TCID_50_).
10. Include mock-infected controls (medium only, no virus) for each experiment to assess background signal.
11. Avoid cross-contamination: use dedicated pipette tips, tubes, and work surfaces for infected samples.
12. Store collected supernatants at -80°C until use. Use aliquots to determine viral titers (e.g., TCID_50_/mL) and viral RNA levels (e.g., RT-qPCR). Fixation timing is critical for preserving antigenicity and tissue morphology. PFUs for infection experiments should be empirically determined for each virus and experimental system (Table 1).
13. Fixation time may be extended to 48 h for thicker or more complex samples in accordance with applicable national biosafety regulations. However, prolonged fixation is associated with antigen masking. As with agarose embedding, homogenization of the fixation and embedding steps in a single continuous process (“single-step casting”) is recommended, as interruptions or partial solidification can create structural weak points. Such discontinuities may lead to fracture lines during cryosectioning. Store frozen blocks at - 80°C until cryosectioning. Avoid repeated freeze-thaw cycles.
14. The temperatures specified here work well for our samples. However, recommended cryostat temperatures depend on the type of tissue and embedding medium, and may vary between cryostats. Always check and determine the appropriate temperature settings for the specific tissue and embedding medium being used before starting sectioning.
15. Proper freezing of the OCT layer is critical to prevent sample slippage during sectioning.
16. After trimming the surrounding OCT to expose the tissue, switch from trim mode to working mode to ensure consistent sectioning. Sections can be cut at 16 µm; thinner sections (e.g., 10 µm) are possible but require optimization of cryostat settings to prevent tearing. If sections break or roll during sectioning, consider adjusting the temperature settings. Also, ensure that the anti-roll plate is clean and properly positioned, and that the blade is sharp (replace the blade if necessary).

## Acknowledgements

LDB, WHI, and SP were supported by the European Union through Horizon Europe (grant agreement 101191666) for the project DEFENDER (‘Identification of novel viral entry factors and development of antiviral approaches’). Part of this work was supported by the BMBF (01KI2058) awarded to TM and SP. SP is additionally associated with the DFG collaborative research center CRC 1648 (SFB 1648/1 2024 – 512741711, ‘Emerging Viruses: Pathogenesis, Structure, Immunity’). We thank the Technology Platform Microscopy and Image Analysis (TP MIA) at the LIV for excellent technical support. We also thank Dr. Aleksandra Kondrashkina (LIV) for her critical review of the method.

